# Postprandial sleep in short-sleeping Mexican cavefish

**DOI:** 10.1101/2024.07.03.602003

**Authors:** Kathryn Gallman, Aakriti Rastogi, Owen North, Morgan O’Gorman, Pierce Hutton, Evan Lloyd, Wes Warren, Johanna E. Kowalko, Erik R. Duboue, Nicolas Rohner, Alex C. Keene

**Affiliations:** Department of Biology, Texas A&M University, College Station, TX 77840; Department of Genomics, University of Missouri, Columbia, MO 65201; Department of Biological Sciences, Lehigh University, Bethlehem, PA 18015; Florida Atlantic University, Jupiter, FL 33458; Stowers Institute for Medical Research, Kansas City, MO 64110

## Abstract

Interaction between sleep and feeding behaviors are critical for adaptive fitness. Diverse species suppress sleep when food is scarce to increase the time spent foraging. Post-prandial sleep, an increase in sleep time following a feeding event, has been documented in vertebrate and invertebrate animals. While interactions between sleep and feeding appear to be highly conserved, the evolution of postprandial sleep in response to changes in food availability remains poorly understood. Multiple populations of the Mexican cavefish, *Astyanax mexicanus,* have independently evolved sleep loss and increased food consumption compared to surface-dwelling fish of the same species, providing the opportunity to investigate the evolution of interactions between sleep and feeding. Here, we investigate effects of feeding on sleep in larval and adult surface fish, and two parallelly evolved cave populations of *A. mexicanus.* Larval surface and cave populations of *A. mexicanus* increase sleep immediately following a meal, providing the first evidence of postprandial sleep in a fish model. The amount of sleep was not correlated to meal size and occurred independently of feeding time. In contrast to larvae, postprandial sleep was not detected in adult surface or cavefish, that can survive for months without food. Together, these findings reveal that postprandial sleep is present in multiple short-sleeping populations of cavefish, suggesting sleep-feeding interactions are retained despite the evolution of sleep loss. These findings raise the possibility that postprandial sleep is critical for energy conservation and survival in larvae that are highly sensitive to food deprivation.

## Introduction

Sleep and metabolic regulation are highly variable throughout the animal kingdom (Lesku *et al*. 2006; Joiner 2016; Keene and Duboue 2018; Seebacher 2018). This variability is reflected by the diversity of food availability and foraging strategy, which potently impact the duration and timing of sleep. There is an interaction between sleep and feeding, regardless of life history strategy, that is critical for organismal survival, and therefore, under selection (Capellini *et al*. 2008; Yurgel *et al*. 2014; Slocumb *et al*. 2015; Aulsebrook *et al*. 2016; Brown *et al*. 2019). While both of these behavioral processes have been studied in detail, much less is known about interactions between sleep and feeding, particularly in the context of evolution.

In many species, sleep deprivation results in increased food intake, while prolonged periods of food deprivation lead to a reduction in metabolic rate and suppression of sleep (Keene *et al*. 2010; Arble *et al*. 2015; Stahl *et al*. 2017; Regalado *et al*. 2017; Goldstein *et al*. 2018). Conversely, animals ranging from the nematode, *C. elegans,* to humans, increase sleep immediately following a meal, revealing an acute effect of dietary nutrients on sleep regulation (Stahl *et al*. 1983; Murphy *et al*. 2016; Makino *et al*. 2021). Defining how evolution has shaped interactions between sleep, metabolic regulation, and feeding is critical to determine the functions of these traits.

The rapidly increasing number of organisms used to study sleep provides new opportunities to study interactions between sleep and metabolism(McNamara *et al*. 2009; Anafi *et al*. 2019). Fish have become a model to study the biological basis of sleep regulation (Chiu and Prober 2013; Levitas-Djerbi and Appelbaum 2017; Keene and Appelbaum 2019). Growing evidence suggests the genetic and functional basis of sleep is conserved across multiple fish species (Chiu and Prober 2013; Levitas-Djerbi and Appelbaum 2017; Keene and Appelbaum 2019). Further, the small size and amenability to genetic manipulation of these fish allows for high-throughput genetic and pharmacological screens to identify novel regulators of sleep (Rihel *et al*. 2010; Chiu *et al*. 2016; Kroll *et al*. 2021). Furthermore, at larval stages, many fish models are transparent, allowing for mapping of sleep and feeding circuits across the entire brain (Semmelhack *et al*. 2014; Leung *et al*. 2019; Wee *et al*. 2019; Förster *et al*. 2020). Therefore, zebrafish and other fish models are exceptionally well positioned to examine interactions between sleep and feeding.

The Mexican tetra, *A. mexicanus* exist as river-dwelling surface fish and at least 30 blind populations of cavefish, which have evolved in nutrient-limited environments, providing the opportunity to examine sleep after fasting and postprandial sleep in an evolutionary context (Jeffery 2009; Gross 2012; McGaugh *et al*. 2020). Multiple cavefish populations have evolved behavioral and physiological differences relative to surface fish including sleep loss, reduced metabolic rate, and increased feeding (Duboué *et al*. 2011; Moran *et al*. 2014; Aspiras *et al*. 2015; Yoshizawa 2015; Volkoff 2016). Long-term starvation has opposing effects on sleep between the surface and cave populations. Starved surface fish suppress sleep, while starved cavefish increase sleep, suggesting that the evolutionary factors shaping the sleep-feeding interaction differ between populations (Jaggard *et al*. 2018). However, sleep-feeding interactions are poorly understood, and postprandial sleep has to our knowledge not been identified in any fish model to date. Examining the effects of feeding state on sleep in surface and cave populations of *A. mexicanus* has the potential to identify whether these behaviors evolved through shared genetic mechanisms and to provide insight into how sleep-feeding interactions are influenced by adaptation to a nutrient-poor cave environment.

Larval *A. mexicanus* provide a particularly tractable model for examining the effects of feeding on sleep regulation. Multiple populations of cavefish larvae have converged on sleep loss similar to adults (Duboué *et al*. 2011; Yoshizawa *et al*. 2015). However, while adult fish can live for months without food, larval fish live for only a matter of days(Salin *et al*. 2010; Medley *et al*. 2022; Pozo-Morales *et al*. 2024). Therefore, interactions between feeding and other behaviors may be particularly important for the survival of larvae and young juvenile fish. Feeding larval fish *Artemia* is readily quantifiable and large numbers of larval fish can be tested without the need to grow fish to adulthood (Espinasa *et al*. 2014, 2017; Lloyd *et al*. 2018). The experimental amenability of larval fish allows for efficient characterization of sleep-feeding interactions across different behavioral and genetic contexts, providing a model to investigate the evolutionary relationship between these processes.

Here, we characterize the effects of starvation and acute feeding on sleep in surface fish and multiple *A. mexicanus* cavefish populations. We identify multiple sleep-feeding interactions in *A. mexicanus*, including the presence of post-prandial sleep in multiple, parallelly evolved cavefish populations. Feeding promotes sleep, independent of time-of-day, revealing the presence of postprandial sleep in both surface and cavefish. Together, these findings reveal interactions between feeding and sleep and provide a model system to examine how these interactions evolved.

## Results

To investigate the effects of feeding on sleep, we compared sleep in different populations of cavefish immediately following a meal. Briefly, fish were fed a meal, and baseline sleep and activity were measured for 24 hours prior to sleep and feeding measurements. At Zeitgeber Time (ZT) 0 on the second day, fish were fed 70 *Artemia* over two hours, followed by a four-hour recording of sleep (Fig 1A). In agreement with previous findings, baseline sleep was lower in both Pachón and Tinaja cavefish compared to surface fish (Fig 1B; Duboué et al. 2011a; Jaggard et al. 2020; O’Gorman et al. 2021a). When sleep was measured following a two-hour feeding period, surface fish slept significantly more than cavefish from both populations (Fig 1C). Consistent with previous findings, quantification of *Artemia* consumed during the two-hour feeding window revealed significantly greater consumption in Tinaja fish, but not Pachón cavefish, compared to surface fish (Aspiras *et al*. 2015; Alié *et al*. 2018)(Fig 1D). Taken together, these findings reveal difference in sleep and feeding behavior of larval *A. mexicanus* populations.

**Figure 1.**
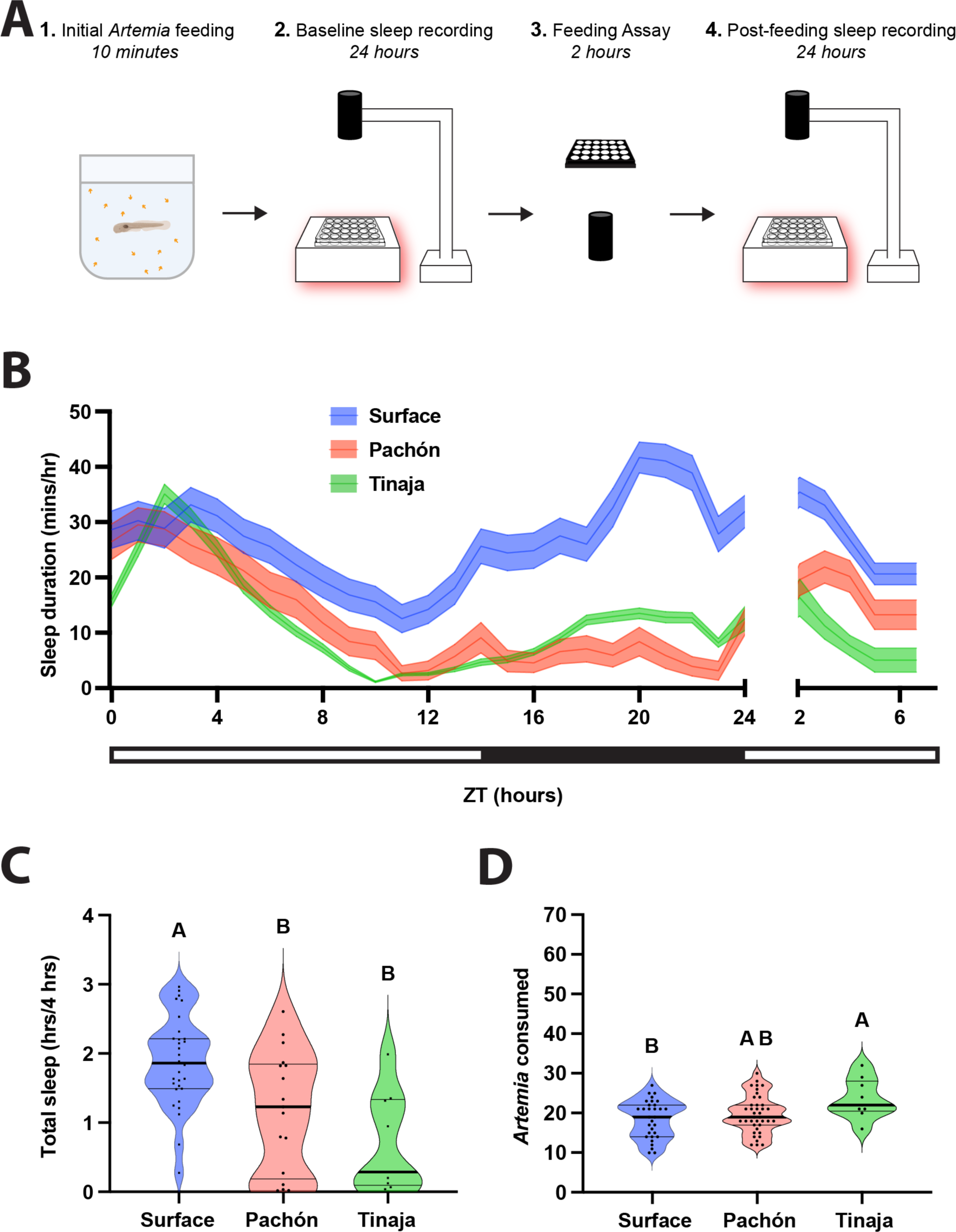
Sleep, feeding, and post-prandial sleep behaviors across three populations of wild-type *Astyanax mexicanus*. **A)** 20 dpf fish were briefly fed prior to 24 h behavioral sleep recordings. At ZT0 the following day, fish were assayed for feeding behavior until ZT2, immediately after which we recorded sleep behaviors between ZT2 and 6. **B)** Sleep profiles of wild type surface, Pachón, and Tinaja fish taken over the experiment. Lines and error bars represent the mean ± SD. **C)** Cross-population comparison of total sleep duration immediately following the feeding experiment. Cavefish slept significantly less than surface fish (ANOVA: F_2, 34_ = 8.123, p = 0.0013; Tukey’s HSD for surface-Pachón, p = 0.0202, p = 0.0024; Tukey’s HSD for surface-Tinaja, p = 0.0024). **D)** Cross-population comparison of the number of *Artemia* eaten during the two-h feeding experiment. Tinaja ate significantly more than surface fish (ANOVA: F_2, 76_ = 3.91, p = 0.0242; Tukey’s HSD for surface-Tinaja, p = 0.0178).

It is possible that sleep is elevated across *A. mexicanus* populations from ZT2-ZT6 due to postprandial sleep or light-regulated rest-activity rhythms. To differentiate between these possibilities, we compared sleep following meals prior to ZT2, ZT6, and ZT10. Feeding time was limited to half an hour to provide additional resolution for postprandial sleep (Fig 2A-C). Across feeding time courses, surface fish slept more than cavefish populations (Fig 2D-F), supporting the notion that surface fish sleep more than cavefish independent of feeding treatment. To measure for postprandial sleep, we compared sleep duration during the four hours following feeding to the remaining hours of daytime (excluding the time for the feeding assay) to determine the percent change in sleep post feeding. Sleep was increased following the meal across all three timepoints, for surface fish and both cavefish populations (Fig 2G-I). Strikingly, for all timepoints tested, there was a significant increase in the amount of postprandial sleep, measured by the increase over the baseline sleep (Fig 2G-I). Variation in the degree of postprandial sleep increase across populations were dependent of feeding time. There were no differences in the percent increase in postprandial sleep between populations fed prior to ZT2, but Surface fish had a significantly greater increase in postprandial sleep than Tinaja cavefish fed prior to ZT6, and Pachón fish had a significantly greater increase in postprandial sleep than either surface and Tinaja cavefish fed prior to ZT10. Similarly, both surface and Pachón cavefish, but not Tinaja cavefish, experienced a significantly greater increase in postprandial sleep prior to ZT10 than for the timepoints earlier in the day. Therefore, while postprandial sleep occurs across *A. mexicanus* populations, the degree to which sleep is increased in each population is dependent on the time of day that feeding occurs. Taken together, these findings reveal the presence of postprandial sleep in surface and cave populations of *A. mexicanus*.

**Figure 2:**
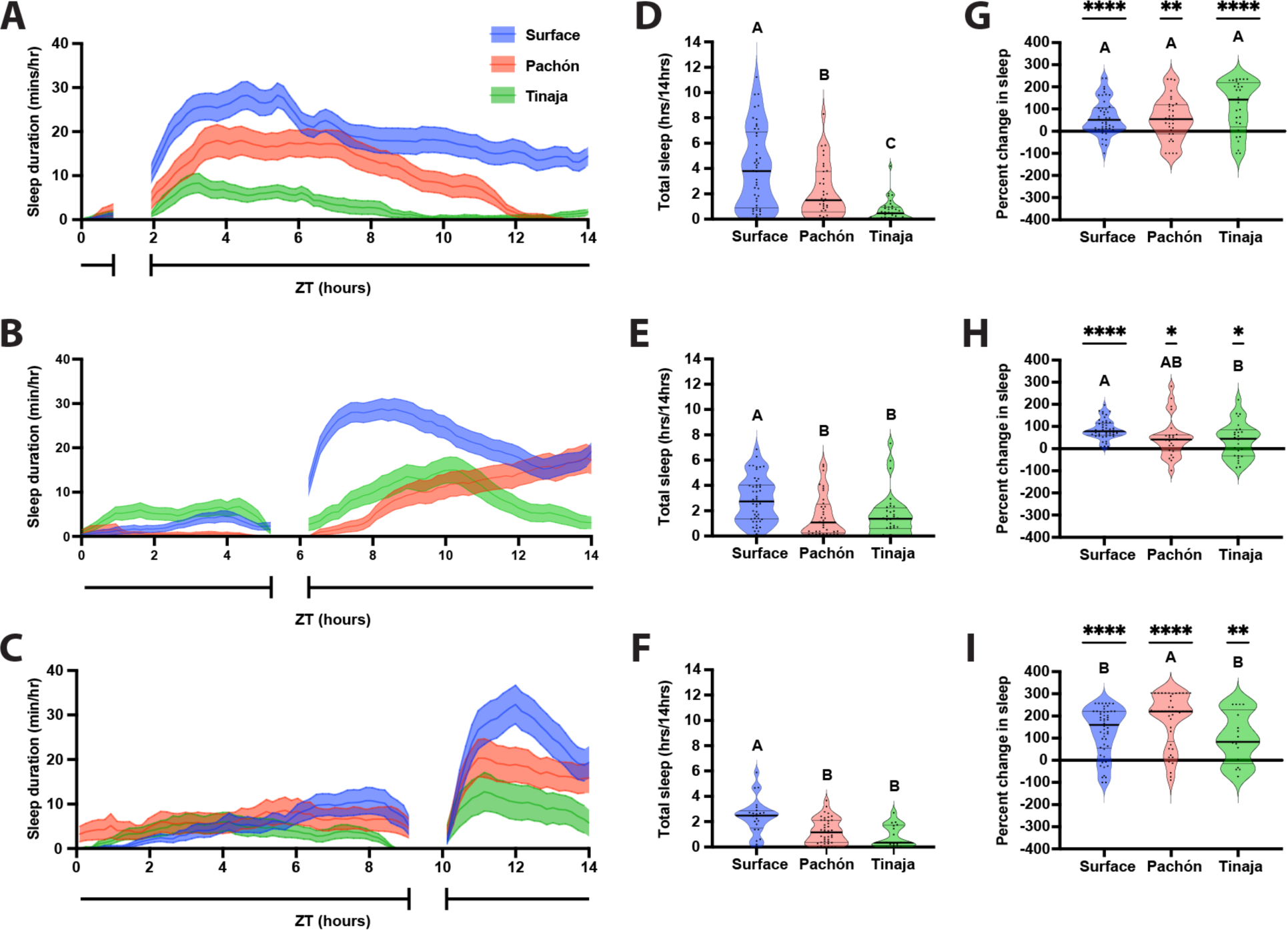
Post feeding increase in larval *A. mexicanus* sleep duration is not dependent on daily feeding time. 20 dpf larvae were fed over a 45-minute window before ZT2 **(A, D, G)**, ZT6 **(B, E, H)**, or ZT10 **(C, F, I)**. **A-C)** Sleep profiles of Surface, Pachón, and Tinaja larvae, in minutes per hour, averaged across the daylight cycle. Lines and error bars represent the mean ± SD. **D, E, F)** Cross-population comparison of total sleep duration in hours over the 14-hour light cycle. Letters represent significant differences. **D)** Total sleep duration around a ZT2 feeding window was significantly different between populations of *A. mexicanus* (ANOVA: F_2, 113_ = 20.81, p < 0.0001). **E)** Total sleep duration around a ZT6 feeding window was significantly different between surface and cave populations of *A. mexicanus* (ANOVA: F_2, 113_ = 8.48, p = 0.0004; Tukey’s HSD for Surface-Pachón, p = 0.001 and Surface-Tinaja, p = 0.0069). **F)** Total sleep duration around a ZT10 feeding window significantly different between surface and cave populations of *A. mexicanus* (ANOVA: F_2, 81_ = 11.64, p < 0.001; Surface-Pachón, p = 0.0003; Tukey’s HSD for surface-Tinaja, p = 0.0002). **G-I)** Percentage change in sleep duration for the four-hour period following feeding from total day time sleep calculated as (proportion of post prandial sleep - proportion of total sleep)/proportion of total sleep. Asterisks indicate significant differences from zero percent change. Letters indicate cross population comparison. **G)** Percent change of postprandial sleep after ZT2 feeding window. Surface: t = 5.333, df = 45, p < 0.0001; Pachón: t = 3.192, df = 31, p = 0.0032; Tinaja: t = 5.239, df = 28, p < 0.0001. There was no significant difference across populations in the percentage of increase in postprandial sleep (Anova: F_2, 104_ = 3.36, p = 0.0417). **H)** Percent change of postprandial sleep after ZT6 feeding window. Surface: t = 13.65, df = 47, p < 0.0001; Pachón: t = 2.67, df = 23, p = 0.0137; Tinaja: t = 2.480, df = 26, p = 0.0200. There was no significant different in the percentage of increase in postprandial sleep between surface and Pachón cavefish, but surface fish had a significantly greater increase in sleep than Tinaja cavefish (ANOVA: F_2, 96_ = 5.758, p = 0.0072; Tukey’s HSD for surface-Tinaja, p = 0.0101). **I)** Percent change of postprandial sleep after ZT10 feeding window. Surface: t = 8.619, df = 52, p < 0.0001; Pachón: t = 10.27, df = 43, p < 0.0001; Tinaja: t = 3.636, df = 16, p = 0.0022. Pachón cavefish had a significantly greater percent increase in postprandial sleep than both surface and Tinaja cavefish (ANOVA: F_2, 111_ = 4.727, p = 0.0107; Tukey’s HSD for surface-Pachón, p = 0.0298; Tukey’s HSD for Pachón-Tinaja, p = 0.0275). For surface fish and Pachón cavefish, the percentage of increase in postprandial sleep was significantly greater after a ZT10 feeding window than at any other timepoint (Surface Anova: F_2, 144_ = 13.84, p < 0.0001; Pachón Anova: F_2, 197_ = 19.56, p < 0.0001). There were no other significant differences in the percent increase for postprandial sleep between timepoints or for Tinaja cavefish (Tinaja Anova: F_2, 70_ = 3.978, p = 0.0231).

It is possible that meal size, or its caloric value, contributes to the duration of postprandial sleep. To determine whether the amount of postprandial sleep is related to meal size, we examined the correlation between the number of *Artemia* consumed and the duration of sleep in the four hours following the meal. For surface fish fed prior to ZT2, there was a significant positive correlation between meal size and post prandial sleep, however there was no significant correlation for surface fish fed prior to ZT6 and ZT10 (Fig 3A-C). For both Pachón (Fig 3D-F) and Tinaja (Fig 3G-H) cavefish, there was no correlation between *Artemia* consumed and postprandial sleep. Therefore, postprandial sleep is largely driven by the presence of a meal and does not appear to be directly linked to meal size.

**Figure 3:**
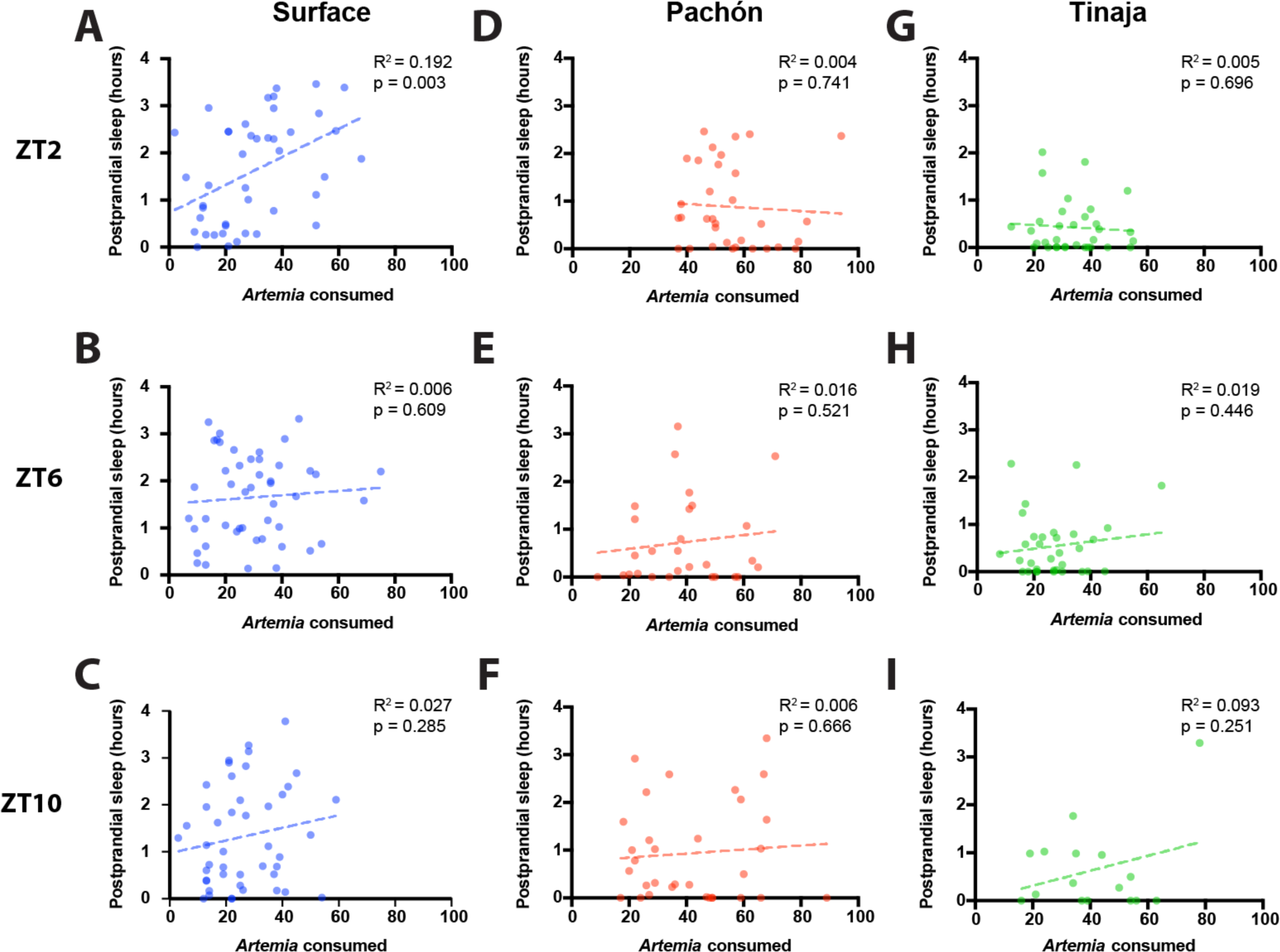
Postprandial sleep in larval *Astyanax* is not dependent on the amount of food consumed, regardless of the time of day that feeding occurs. Correlation of amount of *Artemia nauplii* consumed with sleep duration in the four hours following feeding with a simple linear regression for surface (**A-C)**, Pachón (**D-F)**, and Tinaja (**G-I)**. **A, D, G)** Larvae were fed prior to ZT2. **B, E, H)** Larvae were fed prior to ZT6. **C, F, I)** Larvae were fed prior to ZT10.

Postprandial sleep may provide a mechanism for conserving energy immediately following successful foraging. Conversely, many animals suppress sleep under food-deprived conditions, presumably to forage for food (Macfadyen *et al*. 1973; Danguir and Nicolaidis 1979; Keene *et al*. 2010; Goldstein *et al*. 2018). Larval *A. mexicanus* survive for only a few days without food, raising the possibility that sleep will be acutely impacted by feeding state. To directly examine the effects of feeding state on sleep, we compared sleep in 20 days post fertilization (dpf) fish that were fed from ZT0-ZT2 to unfed fish that had been starved for the previous 24 hours (Fig 4A-C). Surface fish and both populations of cavefish slept significantly more during the four hours following feeding than unfed controls (Fig 4D-F). To further examine the effects of feeding on sleep, we analyzed the activity patterns of fed and unfed fish using a Markov model that predicts the sleep and wake propensity, both indicators of sleep drive (Wiggin *et al*. 2020). Across all three populations, fed fish had a significantly greater sleep propensity P(Doze) and a significantly lower waking propensity P(Wake) than unfed fish, suggesting that sleep drive is increased following feeding (Fig 4G-I). Together, these findings reveal that both surface and cavefish suppress sleep when starved, and that starvation-induced sleep suppression is intact in short-sleeping cavefish.

**Figure 4.**
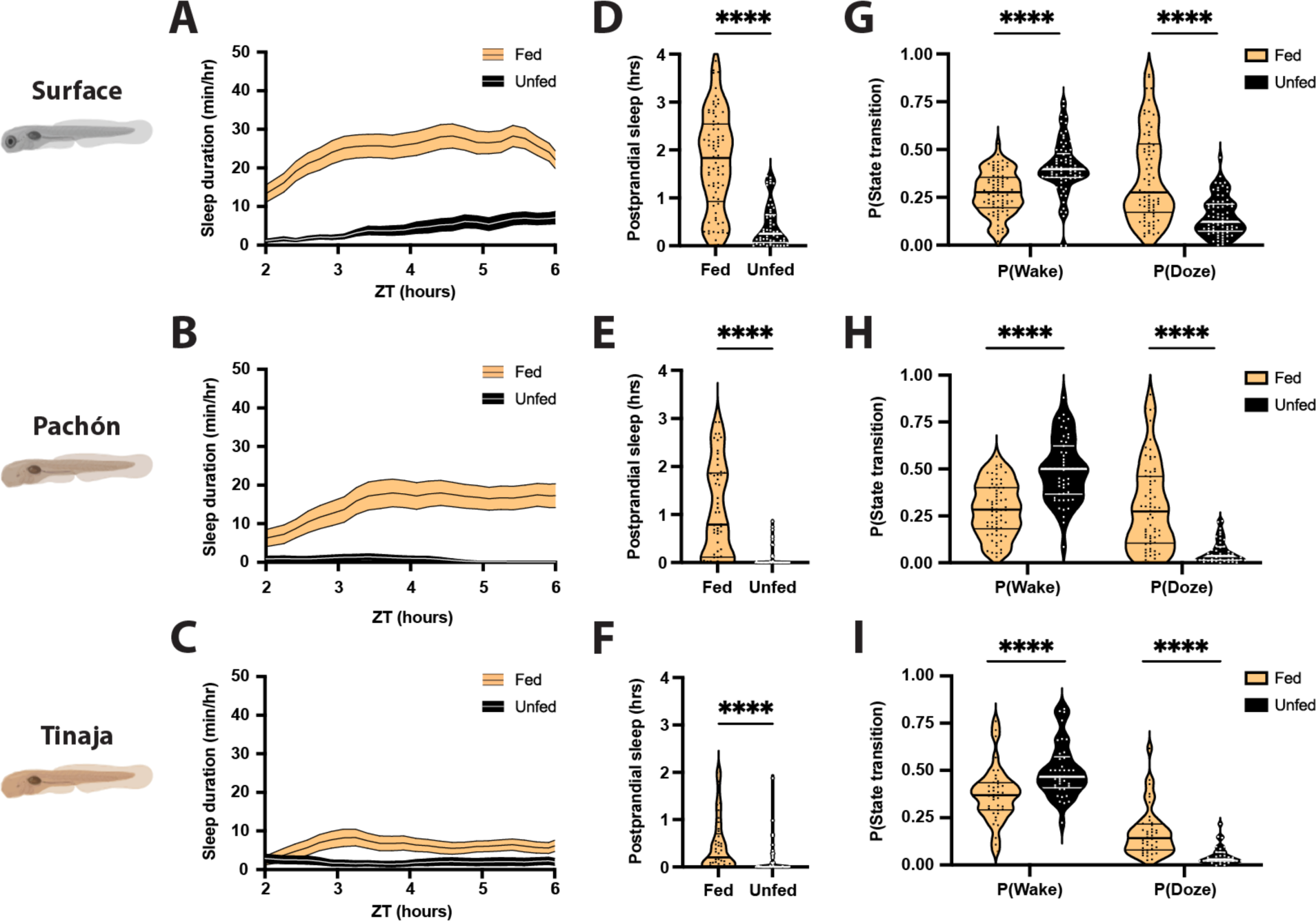
Feeding results in robust increases in sleep duration in larval surface, Pachón, and Tinaja populations of *A. mexicanus*. **A-C)** Four-hour sleep profiles comparing the sleep of fed (orange) and unfed (black) individuals in each population. Lines and error bars represent the mean ± SEM. **D-F)** Fed fish sleep significantly more during the four hours following feeding than unfed fish, regardless of the population. **D)** Surface: Mann-Whitney U = 524, n_fed_ = 77, n_unfed_ = 55, p < 0.0001. **E)** Pachón: Mann-Whitney U = 310.5, n_fed_ = 52, n_unfed_ = 47, p < 0.0001. **F)** Tinaja: Mann-Whitney U = 546.5, n_fed_ = 45, n_unfed_ = 49, p < 0.0001. **G-I)** Fed fish are less likely to wake while asleep, and more likely to fall asleep while awake, than unfed fish. **G)** Surface: P(Wake) Mann-Whitney U = 1317, n_fed_ = 77, n_unfed_ = 76, p < 0.0001; P(Doze) Mann-Whitney U = 1347, n_fed_ = 77, n_unfed_ = 75, p < 0.0001. **H)** Pachon: P(Wake) Mann-Whitney U = 663, n_fed_ = 66, n_unfed_ = 52, p < 0.0001; P(Doze) Mann-Whitney U = 802, n_fed_ = 69, n_unfed_ = 52, p < 0.0001. **I)** Tinaja: P(Wake) Mann-Whitney U = 369, n_fed_ = 40, n_unfed_ = 38, p < 0.0001; P(Doze) Mann-Whitney U = 229, n n_fed_ = 40, n_unfed_ = 34, p < 0.0001. Thin lines represent quartiles.

Adult *A. mexicanus* live months without food and are thought to be highly adapted to survive periods of starvation(Cobham and Rohner 2024). Previously, we have shown that surface fish suppress sleep during periods of prolonged starvation, while cavefish increase sleep (Jaggard *et al*. 2018). To determine whether differences in sleep response extend to acute behavior following meals, we examined postprandial sleep in adult surface and cavefish. Fish were starved for five days prior to recording to synchronize meal patterns and then fed a blood-worm meal at ZT6. In agreement with previous findings(Jaggard *et al*. 2018), control surface fish that were not fed slept significantly more than Pachón and Tinaja cavefish (Fig 5 A, I). Similarly, in fish fed at ZT6, surface fish slept significantly more than Tinaja and Pachòn cavefish (Fig 5B, J). To examine whether postprandial sleep is present in adult *A. mexicanus*, we compared sleep during the four hours following feeding to unfed counterparts (Fig 5C-E). Within this four-hour duration, there were no significant differences in sleep duration (Fig 5F-H) or sleep propensity (Fig 5K-M) between fed and unfed fish across the three *A. mexicanus* populations. Therefore, there is no evident postprandial sleep for adults under the conditions tested, supporting the notion that post prandial sleep is less robust at a life stage when fish are more starvation resistant.

**Figure 5:**
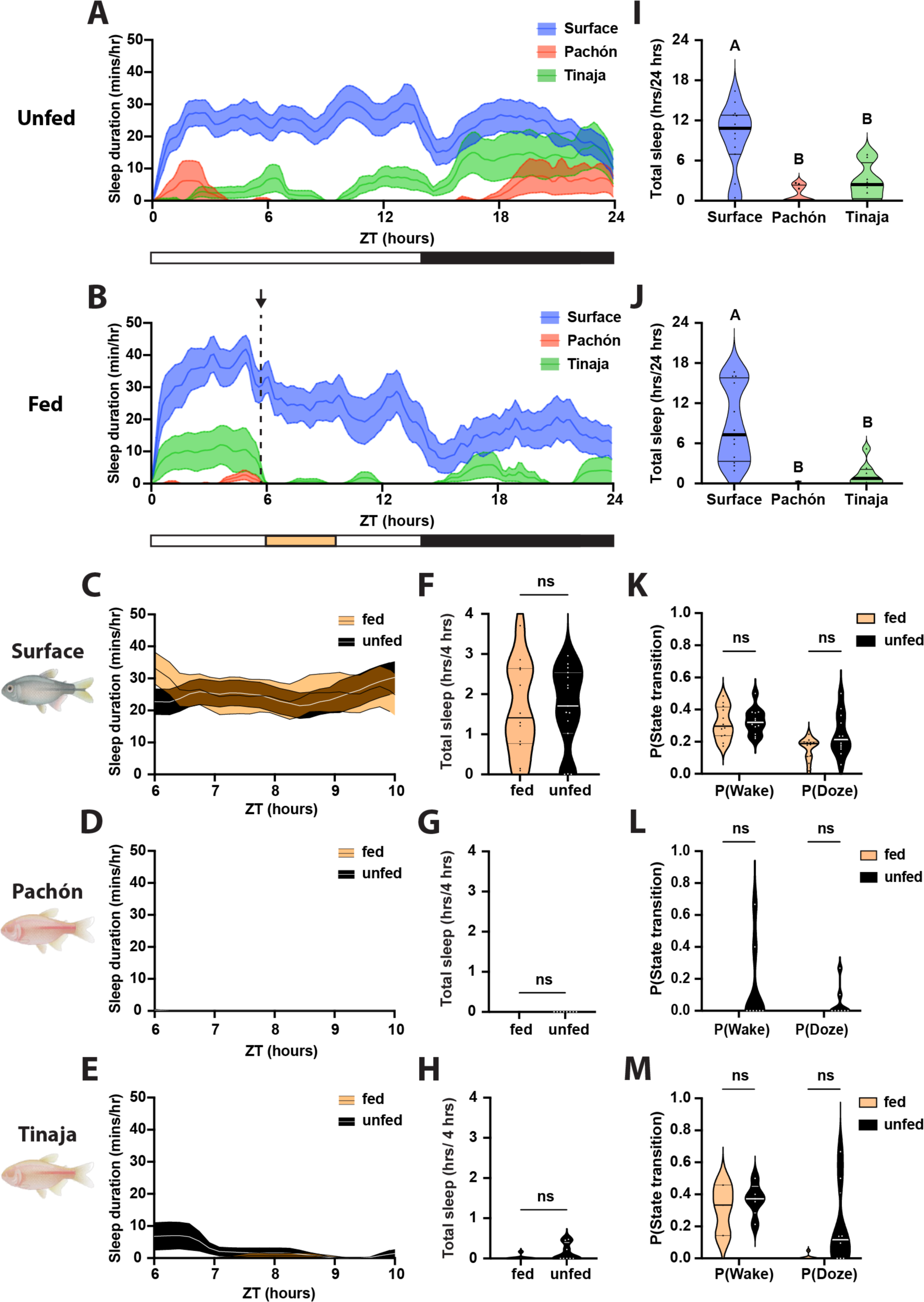
Adult *Astyanax* do not display post prandial sleep behavior. **A, B)** Sleep profiles of adult Surface, Pachón, and Tinaja, in minutes per hour. Lines and error bars represent the mean ± SD. **A, I)** Fish were not fed over the course of the day. **B, J)** Fish were provided food from ZT5.5 (indicated by the arrow and dotted black line in B) to ZT6. **I, J)** Cross-population comparison of total sleep duration in hours over the 24-hour day. Letters represent significant differences. **I)** Total sleep duration in 24 hours was significantly different between unfed surface and cave populations of *A. mexicanus* ((ANOVA: F_2, 28_ = 15.5, p < 0.0001; Tukey’s HSD for Surface-Pachón, p < 0.0001 and Surface-Tinaja, p = 0.0015). **J)** Total sleep duration in was significantly different between fed surface and cave populations of *A. mexicanus* ((ANOVA: F_2, 25_ = 15.04, p < 0.0001; Tukey’s HSD for Surface-Pachón, p < 0.0001 and Surface-Tinaja, p = 0.0008). C-E) Four-hour sleep profiles comparing the sleep of fed (orange) and unfed (black) individuals in each population. Lines and error bars represent the mean ± SEM. **F-H)** There are no significant differences in sleep during the four hours following feeding, regardless of the population. **F)** Surface: Mann-Whitney U = 88, n_fed_ = 12, n_unfed_ = 15, p = 0.9317. **G)** Pachon: Mann-Whitney U = 31.5, n_fed_ = 8, n_unfed_ = 8, p > 0.9999. **H)** Tinaja: Mann-Whitney U = 22.5, n_fed_ = 8, n_unfed_ = 8, p > 0.2. **K-M)** There are no significant differences in activity state transitions between fed and unfed fish. **K)** Surface: P(Wake) t = 0.271, df = 22, p = 0.7888; P(Doze) t = 2.041, df = 22, p = 0.054. **L)** Pachon: Mann-Whitney U = 24, nfed = 8, nunfed = 8; P(Wake) p = 0.4667; P(Doze) p = 0.4667. **M)** Tinaja: Mann-Whitney U = 23, nfed = 8, nunfed = 8; P(Wake) p = 0.5714; P(Doze) p = 0.1319). Horizontal lines represent quartiles.

## Discussion

To date, five populations of *A. mexicanus* cavefish have been studied under laboratory conditions, all of which have significantly reduced sleep compared to surface fish populations (Yoshizawa et al. 2015). These findings have led to the speculation that reduced sleep is adaptive in the food-poor cave environment because it provides more time to forage(Keene *et al*. 2015; Keene and Duboue 2018). However, nearly all studies to date have examined sleep in fed animals, using daily averages. Therefore, little is known about how sleep differs between populations under natural conditions and in response to feeding. Here, we describe interactions between sleep and feeding behavior in surface fish and two different populations of cavefish. All three populations sleep more following feeding than under food-deprived conditions, revealing that feeding is required for baseline sleep. Furthermore, all three populations sleep more in the period following a meal as larvae, but not as adults. These findings suggest that despite robust sleep loss across cavefish populations, sleep-feeding interactions have remained intact.

Numerous neural mechanisms associated with sleep loss in cavefish have been identified including elevated levels of the wake-promoting neuropeptide Hypocretin (HCRT), changes in wake-promoting catecholamine systems (Duboué *et al*. 2012; Bilandzija *et al*. 2013; Gallman *et al*. 2019) providing candidate regulators of postprandial sleep. Similarly, feeding is increased in multiple populations of adult *A. mexicanus* (Aspiras *et al*. 2015). In agreement with previous findings, we find that feeding is elevated in 20 days post fertilization juvenile cavefish from the Tinaja, but not Pachón population (O’Gorman et al. 2021). In adults, differences in feeding are at least partially attributable to polymorphisms in the GPCR Melanocortin 4 receptor (Mc4r) which is associated with obesity in humans and animal models (Aspiras *et al*. 2015). While there is little evidence that MC4R directly regulates sleep, it is thought to contribute to obesity-induced sleep apnea that in turn regulates sleep (Larkin *et al*. 2010; Pillai *et al*. 2014). Our findings that post-prandial sleep is intact in Tinaja cavefish suggests that Mc4r, and other genes involved in feeding, are likely dispensable for sleep feeding interactions. There are also numerous genes that have been identified to regulate sleep or feeding in fish models that are potential regulators of sleep-metabolism interactions. For example, the orexigenic neuropeptides Neuropetide Y (Npy) and Hcrt both induce wakefulness, providing a potential molecular mechanism for feeding-dependent modulation of sleep (Appelbaum *et al*. 2009; Penney and Volkoff 2014; Singh *et al*. 2015, 2017; Jaggard *et al*. 2018). Future functional analysis is required to define whether these candidate genes regulate interactions between sleep and feeding.

In *A. mexicanus*, rhythmic transcription is significantly diminished under dark-dark conditions, and cavefish have elevated levels of light-inducible genes(Beale *et al*. 2013). The circadian clock plays a critical role in the timing of both sleep and feeding, raising the possibility that the circadian clock may be critical for sleep-feeding interactions. Transcriptome-wide analysis in larvae, reveals a loss of rhythmic gene expression across all cave populations tested (Mack *et al*. 2021) Therefore, because identified postprandial sleep in all of the populations tested across three different timepoints during the day, postprandial sleep may be independent of time-of-day and may not require a functioning circadian clock.

*A. mexicanus* larvae, like zebrafish, can subsist on a variety of foods including paramecium, rotifers, and fish feed that differ in micronutrients. In this study, *A. mexicanus* larvae were fed a standard diet of *Artemia. Artemia* is comprised of macronutrients that include diverse fatty acids, proteins, and carbohydrates. Analysis suggests that *Artemia is* ∼40-60% protein, raising the possibility that consumption of dietary protein may impact sleep (de Clercq *et al*. 2005). In *Drosophila,* dietary protein promotes post-prandial sleep, while a loss of dietary protein disrupts sleep depth (Murphy et al. 2016; Brown et al. 2020; Titos et al., 2023). Therefore, it is possible that changes in protein detection, or its downstream targets, regulate the physiology of sleep circuits that are responsible for the different effects of feeding on sleep between Pachón and Tinaja cavefish. Understanding the effects of different diets on sleep, and how individual macronutrients regulate sleep across populations could reveal evolved differences in sleep-feeding interactions across different *A. mexicanus* populations.

The identification of postprandial sleep in cavefish provides an avenue for future studies examining the genetic basis of this behavior. Mapping genetic loci associated with trait variation has been used to identify candidate regulators of many morphological and behavioral traits, including regulators of sleep, activity, feeding posture, and metabolism (Kowalko *et al*. 2013; Yoshizawa *et al*. 2015; Carlson *et al*. 2018; Riddle *et al*. 2021). Further, population genetic approaches have identified genome-wide markers of selection across multiple cave populations, and this genetic variation may provide insight into genes impacting sleep-feeding interactions (Herman et al. 2018; Warren et al. 2021; Moran et al. 2022). Genes with signatures of selection that have previously been implicated in sleep or feeding could provide candidate regulators of postprandial sleep. In *A. mexicanus*, like zebrafish, CRISPR-based gene editing has been used to functionally validate genes identified through genomics approaches and could be applied to the investigation of postprandial sleep (Klaassen *et al*. 2018; Kroll *et al*. 2021). Genetic studies will require the use of CRISPR for forward genetic screens, or the identification of *A. mexicanus* with diminished or highly variable post-prandial sleep that can be used for genetic mapping studies.

In conclusion, these studies identify postprandial sleep in *A. mexicanus* and suggest it is under independent genetic regulation from total sleep duration and meal size in surface fish and two parallely evolved populations of cavefish. These studies lay the groundwork for future analysis that apply currently available population genetics, neural anatomical, and genetic screening toolsets in *A. mexicanus* to examine the integration of feeding and sleep regulation

## Materials and Methods

### Methods

#### Husbandry

Throughout this study, we followed previously described standard animal husbandry and breeding for *A. mexicanus* (Borowsky 2008a). All fish were housed under standard temperature (23°C for adults, 25°C for embryos and larvae) and lighting conditions (14:10 hr light:dark cycle). Adult fish were bred by increasing water temperature to 27±1°C and feeding a high-calorie diet that includes thawed frozen bloodworms three times per day (Elipot *et al*. 2014). Larvae were fed brine shrimp (*Artemia nauplii*) *ad libitum* from 6 – 20 days post-fertilization (dpf; Borowsky 2008b). Embryos and larvae were held in small glass bowls until behavioral testing. All procedures in this study were approved under the Florida Atlantic University and Texas A&M University IACUC.

#### Sleep behavior

These experiments focused on three distinct *A. mexicanus* morphotypes: the sighted, surface-dwelling Río Choy, and two blind, cave-dwelling populations, Pachón and Tinaja. We quantified sleep behavior in these fish using previously described methods (Jaggard *et al*. 2019a) and baseline sleep data (O’Gorman *et al*. 2021). Briefly, we used Ethovision XT 17.0 software (Noldus Information Technology, Wageningen, the Netherlands) to track locomotor behavior. Raw locomotor behavior was used to calculate sleep behavior parameters using a custom Perl script(Jaggard *et al*. 2019b). We operationally define sleep as 60 seconds or more of immobility given that previous studies show both surface and Pachón cavefish exhibit increased arousal thresholds after this period(Jaggard *et al*. 2019b). We defined immobility as a velocity below 6 mm/sec for larval fish and a velocity below 4 cm/sec for adult fish. All recordings were performed at 23 °C under a 14:10 hour light/dark cycle.

#### Larval behavior recordings

All larval used to quantify sleep behavior were 20 dpf. Fish were fed and then acclimated individually in 24-well plates for at least 15 hours prior to behavior recordings. Recordings began at ZT0 and lasted for 24 hours, with interruptions for feeding at specific time points. The 24-well plates were placed on light boxes made from white acrylic housing infrared (IR) lights (Figure 1A). Basler ace acA1300-200um Monochrome USB 3.0 Cameras with mounted IR filters were mounted above the well plates and recordings were taken using Pylon Viewer software.

The effects of feeding on sleep were tested throughout the light cycle at time points prior to ZT0, ZT2, ZT6, and ZT10. Each 24-well plate was either not fed as a control or fed at a single time point. We conducted two separate feeding experiments. In the first experiment, larvae were fed for 10 mins immediately before a 24-hour recording beginning at ZT0. This 24-hour recording was followed by a 2-hour feeding behavior assay (described below) and then another behavior recording for 4 hours from ZT2-ZT6 (Fig 1). In the second experiment, we recorded behavior for 24 hours around a 45-minute window for feeding prior to either ZT2, ZT6, or ZT10.

#### Larval feeding behavior assay

To quantify the relationship between the amount of food consumption and post-prandial sleep duration, we performed feeding assays that allowed us to count the number of *Artemia* over a given time. The duration of the feeding assay was 2 hours for the first experiment, starting at ZT0 following 24 hours of recording. The duration of the feeding assay was 30 minutes for the second experiment, starting prior to ZT2, ZT6, or ZT10. For the 2-hour feeding assay, fish were given exactly 70 *Artemia*, for the 30 minute feeding assay, *Artemia* were provided *ad libitum*. We filled a new 24-well plate with *Artemia* hatched within 24 hours and recorded for at least one minute prior to transferring the larval fish from the recording well plate to this new feeding well plate. At the end of the recording duration, fish were removed from the feeding assay, placed back into the original 24-well recording plate with clean water and returned to the behavior recording. We used FIJI (Schindelin *et al*. 2012) to count the number of *Artemia* both before the fish were added to the wells and at the end of the feeding assay. Subtraction of the former from the latter allowed us to determine the amount of *Artemia* eaten over the duration of the feeding assay.

#### Adult behavior recordings

Adult fish used for behavior recordings were approximately 1 year old with an equal number of males and females per treatment. Food was withheld for 5 days prior to recording. Fish were placed in individual glass tanks of approximately 30 x 17 cm in a 2 x 2 grid in front of an IR light board and left to acclimate for at least 24 hours. Recordings began at ZT0 and lasted 24 hours. In the top two tanks, 4 oz of thawed, frozen blood worms were added at ZT5.5 and any uneaten worms were removed after 30 minutes at ZT6. The fish in the bottom two tanks were not fed as a control.

#### Analysis

Statistical analyses were performed in GraphPad Prism (version # 9.5.0) and R (version 4.0.4). When assumptions of normality and equal variances were met, we used parametric t-tests, ANOVA, and Pearson’s r tests, otherwise we used non-parametric Mann-Whitney U, Kruskal-Wallis, and Spearman’s ρ tests. Following a significant ANOVA or Kruskal-Wallis test, pairwise comparisons were made using Tukey’s HSD or Dunn’s test, respectively.

To quantify the percent change in sleep duration during the 4 hours following feeding, we determined the proportion of total daylight sleep to total daylight recording time as well as the proportion of sleep to the 4 hour post prandial recording period. We then calculated percent change as the proportion of post prandial sleep minus the proportion of total daylight sleep divided by the proportion of total daylight sleep. Finally, to test whether the amount of *Artemia* consumed was related to post-prandial sleep duration, we analyzed the goodness of fit from a linear regression.

## Acknowledgements

This work was supported by NIH Grants NIH 1R01GM127872 to SEM, NR, and ACK; R24 OD030214 to WW, NR and ACK; R21 NS122166 to ACK and JEK; 1DP2AG071466-01 to NR; NSF Grant NSF grant IOS 2202359 to JEK and SEM. The authors are grateful for technical assistance from Kaya Harper and Lawaal Agboola.

